# A structural and functional analysis of opal stop codon translational readthrough during Chikungunya Virus replication

**DOI:** 10.1101/2023.07.24.550286

**Authors:** Raymond Li, Kaiwen Sun, Andrew Tuplin, Mark Harris

**Affiliations:** School of Molecular and Cellular Biology, Faculty of Biological Sciences, and Astbury Centre for Structural Molecular Biology, University of Leeds, Leeds, West Yorkshire, United Kingdom LS2 9JT

## Abstract

Chikungunya virus (CHIKV) is an alphavirus, transmitted by *Aedes* species mosquitoes. The CHIKV positive-sense RNA genome contains two open reading frames, coding for the non-structural (nsP) and structural proteins of the virus. The non-structural polyprotein precursor is proteolytically cleaved to generate nsP1-4. Intriguingly most isolates of CHIKV (and other alphaviruses) possess an opal stop codon close to the 3’ end of the nsP3 coding sequence and translational readthrough is necessary to produce full-length nsP3 and the nsP4 RNA polymerase. Here we investigate the role of this stop codon by replacing the arginine codon with all three stop codons in the context of both a subgenomic replicon and infectious CHIKV. Both opal and amber stop codons were tolerated in mammalian cells, but the ochre was not. In mosquito cells all three stop codons were tolerated. Using SHAPE analysis we interrogated the structure of a putative stem loop 3’ of the stop codon and used mutagenesis to probe the importance of a short base-paired region at the base of this structure. Our data reveal that this stem is not required for stop codon translational readthrough, and we conclude that other factors must facilitate this process to permit productive CHIKV replication.

## Introduction

Chikungunya virus (CHIKV) is a positive-sense single stranded RNA virus, a member of the Alphavirus genus in the Togaviridae family which infects both mosquitoes and mammalian hosts. Alphavirus infections in mosquitoes are life-long and symptomless but infection in mammalian hosts causes acute infection. The main vectors for alphavirus infection are *Aedes aegypti* and *Ae. albopictus* mosquitoes, and the increasing global distribution of alphaviruses has been attributed to climate change resulting in an increase in the vector range, as well as increased movement of people via tourism and migration. CHIKV was first isolated in Tanzania in 1952. There are three strains of the virus: the East/Central and South African (ECSA), the West African and the Asian strains. Infection with CHIKV causes a non-fatal febrile illness in infected humans that normally lasts for months after the initial infection. However, CHIKV can progress to chronic infection in 40% of cases, and individuals suffer from long term painful persistent joint and muscle pain [1]. There are no licensed vaccines or specific antivirals available for CHIKV, current treatment consists of nonsteroidal anti-inflammatory drugs to provide symptom relief.

The CHIKV genome is a capped 11.8 kb RNA with a 3′ poly-(A) tail comprising two open reading frames (ORFs) and three untranslated regions (UTRs) located at the 5′ and 3′ ends, and between the two ORFs. The first ORF is translated into the non-structural proteins nsP1 to nsP4. The second ORF is translated into the structural proteins (capsid, glycoproteins E1 to E3, and the 6K protein and is encoded on a subgenomic (sg) RNA, transcribed from a SG promoter [2].

The nsPs perform essential functions during viral replication, nsP1 caps the 5’ end of the genome and sgRNA [3], nsP2 is a protease, nucleoside triphosphatase and helicase [4] and nsP4 is the RNA dependent RNA polymerase (RdRp) [5]. The exact functions of nsP3 are currently poorly characterised. The viral protein has roles during the early stages of infection as nsP3 is necessary for synthesis of the negative sense RNA intermediate [6] with other studies revealing importance for the transcription of sgRNA [7]. NsP3 comprises three domains: The N-terminal macro domain, central alphavirus unique domain (AUD) and C-terminal hypervariable domain (HVD).

Intriguingly, an opal stop codon (UGA) has been identified in nsP3, 6 codons before the cleavage site between nsP3 and nsP4, in some isolates of CHIKV and other alphaviruses (Fig. 1a). In other CHIKV isolates however, this opal stop codon is substituted by an arginine codon (AGA or CGA). The reason for the presence or absence of the opal stop codon is poorly understood [8], however there must be significant translational read-through (TR) of this stop codon otherwise the nsP4 RdRp could not be produced and the virus would be unable to undergo genome replication [9]. In addition, different isoforms of nsP3 are expressed when either TR or termination at the opal stop codon occurs, the exact function of the two forms of nsP3 and the role during the virus lifecycle are currently unknown. In the case of the prototypic alphaviruses Semliki Forest Virus (SFV) and Sindbis virus (SINV) the 6 C-terminal residues of SFV nsP3 were shown to promote protein degradation reducing the half-life of nsP3 8-fold [10]. It is possible that this degradation signal is conserved amongst alphaviruses, the opal stop codon TR mechanism may function to drive expression of two populations of nsP3 which function in different aspects of the virus lifecycle. In the case of CHIKV, a Caribbean isolate was shown to be a quasispecies where both the arginine and opal stop codon were detected in different virus samples. Replacing the opal stop codon of a Sri Lankan strain of CHIKV by an arginine codon did not affect virus replication *in vitro* or *in vivo* but caused decreased pathogenesis in a mouse model. Presence of the arginine codon in the virus caused a delayed proinflammatory chemokine and cytokine recruitment phenotype resulting in reduced ankle swelling compared to the opal stop codon mutant [11].

**Figure 1.**
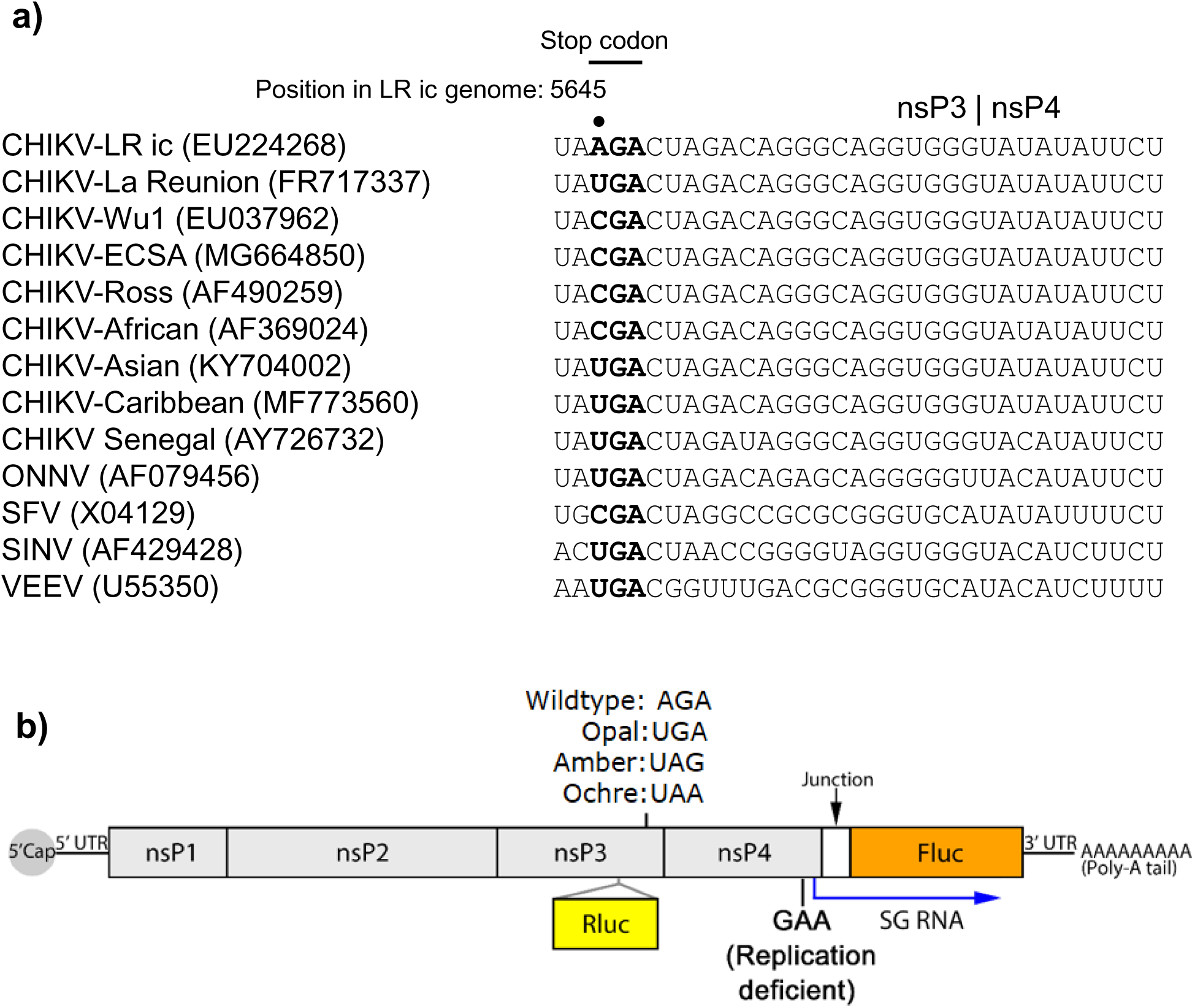
(a) An alignment of the nucleotide sequences at the stop codon and 3’ illustrating the conservation of sequence at the end of the nsP3 coding region. (b) Schematic of the CHIKV-Dluc-SGR used in this study. Renilla luciferase (Rluc) is expressed as an internal fusion with nsP3, firefly luciferase (Fluc) translated from a subgenomic RNA.

TR has been reported to be stimulated by the presence of a 3’ located RNA structure [12]. In this context a recent study used selective 2′ hydroxyl acylation and primer extension (SHAPE) to experimentally determine RNA structures in short sequences of the African/Asian or Caribbean strains of CHIKV inserted into bicistronic reporter plasmids [13]. These data confirmed the presence of a large RNA structure and furthermore demonstrated that it functioned to promote opal stop codon TR [13]. However, it is currently unknown whether this RNA structure could stably exist at lower temperatures (eg in the mosquito vector), or indeed whether it exists in the context of the full length CHIKV genome. We therefore sought to investigate the influence of stop codon TR on CHIKV replication in mammalian and mosquito cells. To do this the opal, ochre and amber stop codon were introduced into an ECSA strain of CHIKV which encodes an arginine at this position in the context of an SG replicon with dual luciferase (Dluc) reporter proteins (Fig. 1b). These constructs were screened in a panel of cell types to investigate cell type specific phenotypes for the stop codon read-through mutants and the mutations were then investigated in the context of a full infectious clone of the ECSA CHIKV strain. The second objective was to provide further insight into the mechanism of stop codon TR for CHIKV in the context of the mammalian host and mosquito vector. This was accomplished by experimentally determining the structure of the conserved stop codon TR RNA structure in the context of the full infectious genome folded at temperatures corresponding with the human and mosquito body temperature. We also show that a conserved base-paired region at the base of the stem is not required for virus replication in mammalian cells.

## Results

### Effect of stop codons in nsP3 on CHIKV genome replication analysed using a CHIKV subgenomic replicon

In many alphaviruses an opal stop codon is located 6 codons upstream of the nsP3-nsP4 cleavage site, the presence of the stop codon results in the expression of the nsP123 precursor as the major polyprotein product. TR at the stop codon occurs approximately 10% of the time to yield the full nsP1234 polyprotein [14]. Subsequent cleavage of the nsP1234 polyprotein yield the active RdRp nsP4 and thus read-through of the stop codon is essential for virus genome replication. The presence of a stop codon is thus enigmatic and as yet unexplained. Intriguingly, isolates of CHIKV have been identified with either an opal stop codon (UGA) or a sense codon (arginine: AGA or CGA) at this position (Fig. 1a). The ICRES isolate of CHIKV (LR2006 OPY1) [15] has been used extensively by many groups including ourselves to investigate aspects of CHIKV biology and contains an arginine codon (Fig. 1a). To investigate the effect of a stop codon at this position on CHIKV genome replication, we first replaced the arginine codon with each of the three stop codons in the context of a CHIKV subgenomic replicon (SGR) in which the structural protein encoding region was replaced with firefly luciferase (Fluc), and renilla luciferase was expressed as an internal fusion with nsP3 (Fig. 1b: CHIKV-Dluc-SGR). The wildtype arginine (AGA) codon (residues 5645-47) was substituted with the opal (UGA), ochre (UAA) or amber (UAG) stop codons. As a negative control we generated a polymerase inactive nsP4 mutant in which the active site was mutated (GDD → GAA). In this SGR, Rluc expression reflects both input RNA translation and early genome replication whereas Fluc expression is dependent on genome replication and transcription. Fluc expression is thus also absolutely dependent on successful stop codon TR; the stop codon is before the nsP4 encoding sequence and thus early termination would prevent the expression of the nsP4 RdRp required for active genome replication complex.

We first investigated the phenotype of the stop codon derivatives of CHIKV-Dluc-SGR in a variety of mammalian cells, identified previously as efficiently supporting CHIKV genome replication [16]. Specifically we used human hepatocellular carcinoma cell line Huh7, the human glial cell line SVG-A and the human muscle derived rhabdomyosarcoma cell line RD, representing target cell types for CHIKV replication in infected humans [17]. As shown in Fig. 2 (a-c), wildtype CHIKV-Dluc-SGR replicated efficiently in all three cell lines with Fluc values displaying a ∼1000-fold increase between 4-24 h post transfection (hpt), consistent with previous observations [16]. The presence of the opal stop codon had no significant effect on replication in Huh7 cells but displayed a modest, statistically significant, reduction in replication in SVG-A and RD cells. In contrast, the presence of the amber stop codon resulted in a 10-fold reduction in replication in all 3 cell types. The presence of the ochre stop codon had a more dramatic effect: a ∼100-fold decrease in Huh7 cells but was indistinguishable from the negative control (nsP4 GAA) in SVG-A and RD cells. These phenotypes were also reproduced in murine myoblast cells, C2C12 (Fig. 2d), previously shown to support very high levels of CHIKV replication [16]. We conclude that in mammalian cells efficient TR at both the opal and amber stop codons occurs, however the ochre stop codon cannot undergo TR.

**Figure 2.**
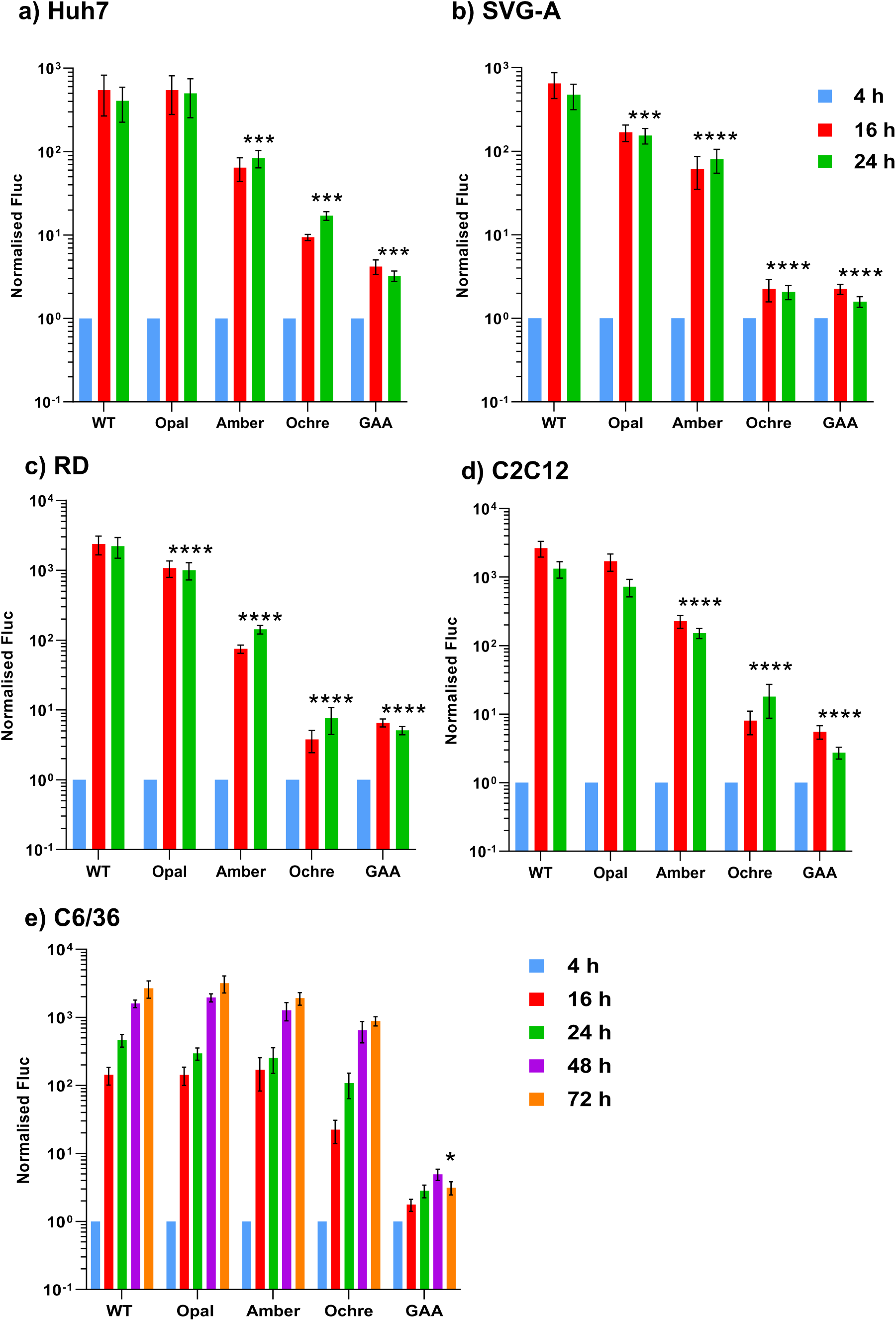
Replication of CHIKV-Dluc-SGR wildtype and stop codon mutants in different cell lines. Graphs show Fluc values normalised to the 4 h post transfection value.

As CHIKV is transmitted via a mosquito vector we also assessed the phenotype of the mutant panel in C6/36 cells, derived from the mosquito *Aedes albopictus*. Surprisingly, as shown in Fig. 2e, replication of all three stop codon mutants was indistinguishable from wildtype in these cells suggesting that efficient TR at all three stop codons occurred in mosquito cells.

### Effect of stop codons in nsP3 on production of infectious virus

We proceeded to investigate the phenotype of the stop codons in the context of the complete viral lifecycle. We reasoned that reduced translation of the nsP4 RdRp in the absence of TR would limit transcription of the subgenomic RNA encoding for the structural proteins (Fig. 3a), thus reducing production of infectious virus. The stop codons were therefore introduced into the ICRES infectious clone [15], from which the SGR had been derived. Virus stocks were produced by transfection of *in vitro* transcribed RNA into BHK-21 cells as these have been shown to be highly permissive for alphavirus replication [18]. The transfected cells were first analysed by infectious centre assays (ICA) to determine if the presence of the stop codons had an effect on the production of infectious CHIKV particles. The ICA protocol directly determines the production of infectious virus from the transfected RNA, circumventing any possibility of reversion. The ICA involved serially diluting transfected cells and overlaying them on to a monolayer of naïve cells. Plaques observed in the monolayer corresponded to transfected cells releasing infectious virus to neighbouring cells. Consistent with the replication data the opal stop codon mutant exhibited the same RNA infectivity (plaque forming units per µg RNA) as the wildtype and the amber stop codon resulted in a ∼ 6-fold reduction (Fig. 3b). In contrast the ochre stop codon completely abrogated RNA infectivity. The same results were observed when supernatants harvested at 24 hpt were titrated by plaque assay (Fig. 3c), the absence of reversion was confirmed by RT-PCR and sequencing analysis of RNA extracted from the transfected cells at 24 hpt (data not shown). Note that no RT-PCR product could be generated from the ochre stop mutant electroporated cells, consistent with the absence of infectious virus by ICA or plaque assay. We also investigated the phenotype of the infectious clone stop codon mutants in C6/36 cells, in this experiment plaque assays were performed with 48 hpt supernatants to reflect the slower rate of replication in these cells (see Fig. 2). Consistent with the SGR data both the opal and amber stop codons exhibited a similar infectivity to wildtype (Fig. 3d), although intriguingly in this case the ochre stop codon resulted in a 1000-fold reduction. Again the absence of reversion was confirmed by RT-PCR and Sanger sequencing analysis of RNA extracted from the transfected cells at 48 hpt (data not shown).

**Figure 3.**
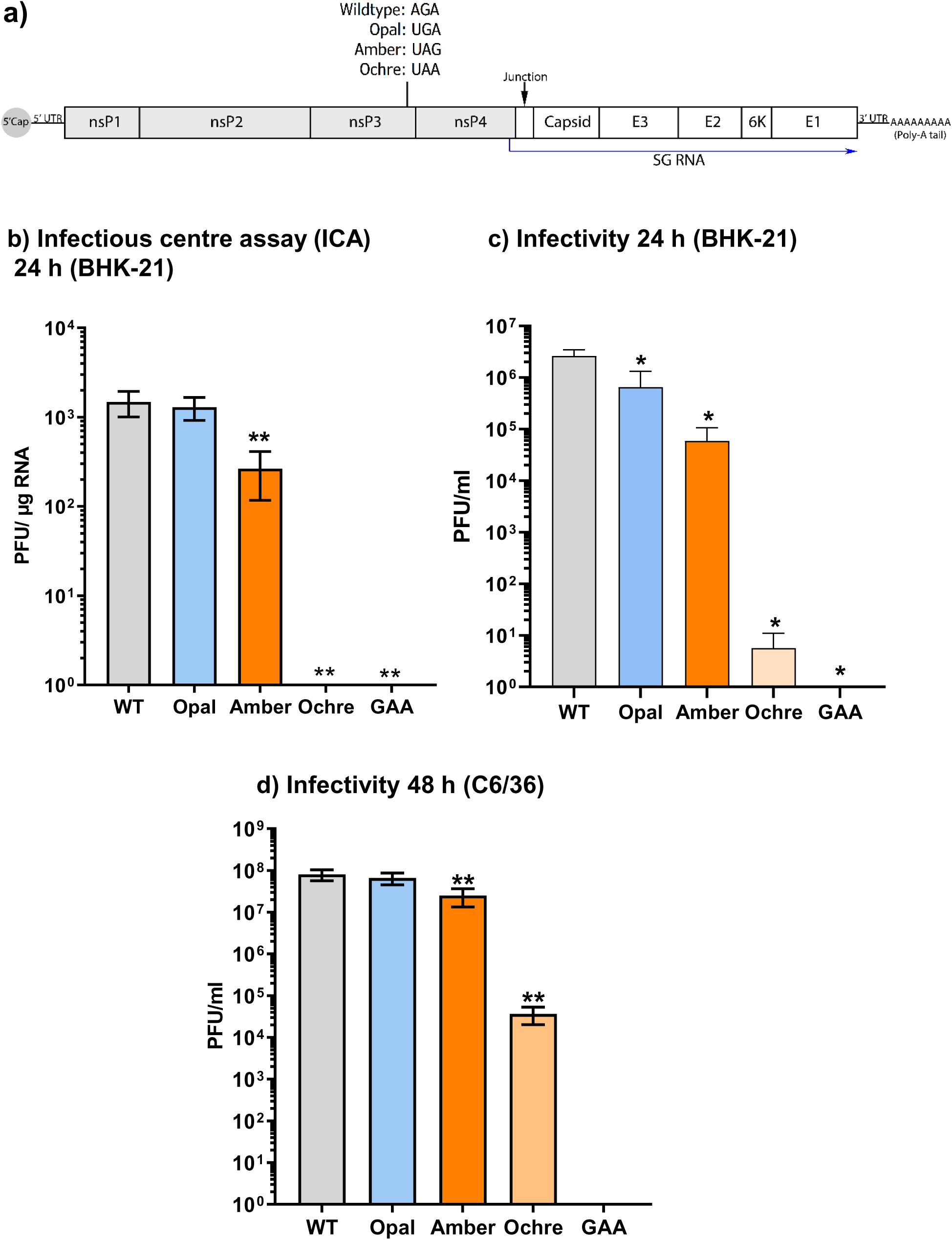
(a) Schematic of the CHIKV infectious genome showing location of the stop codons. (b) Infectious centre assay (ICA) for BHK-21 cells electroporated with *in vitro* transcribed CHIKV RNA at 24 h post electroporation. Graph shows number of plaques formed in a BHK-21 monolayer overlaid with electroporated cells expressed relative to the µg of RNA electroporated. (c) Plaque assay of virus released from electroporated BHK-21 cells at 24 h, expressed as plaque forming units per ml of clarified supernatant (PFU/ml). (d) Plaque assay on BHK-21 cells of virus released from electroporated C6/36 cells at 48 h, expressed as PFU/ml of clarified supernatant.

### Structural analysis of a downstream element previously shown to be required for stop codon TR

The exact mechanism of alphavirus stop codon TR is currently unclear, however previous work has identified two contributory elements. Firstly, mutagenesis of a cytidine base immediately 3’ of the stop codon disrupted TR of the SINV opal stop codon [14]. Secondly, a highly conserved RNA structure located within the 150 nucleotides 3’ to the stop codon [12]. A stable stem at the base of this structure was shown to be important for TR in the context of a plasmid reporter construct [12]. More recently this structure from CHIKV was mapped by SHAPE analysis [13] in the context of a short RNA fragment derived from the CHIKV genome.

We sought to further analyse the structure of this element using QuSHAPE, which utilises fluorescently labelled primers and is analysed by capillary electrophoresis analysis [19] to provide quantitative values of NMIA reactivity at a single nucleotide resolution. Given the different phenotypes of the stop codon mutants in mammalian and mosquito derived cells we hypothesised that the overall structure of this element might be dependent on the temperature. We therefore performed the QuSHAPE analysis using full-length ICRES CHIKV genome RNA folded at 37°C and 28°C. We considered that QuSHAPE experiments on the full-length CHIKV RNA would reveal whether other parts of the genome could influence the formation of the read-through RNA structure.

In fact, the averaged NMIA reactivity for the 37°C and 28°C folded RNA produced relatively similar profiles across the element (Fig. 5a). Predicted RNA structure models generated using the RNAstructure software [20] constrained by the experimental SHAPE data for the 37°C (Fig. 5b) and 28°C (Fig. 5c) folded RNA confirmed a similar large RNA structure 3′ to the opal stop codon position. The reactivity between bases 5694 to 5769 form peaks which are highly similar in size and shape. Some changes are seen for the 28°C reactivity for the 5′ stem which appears to be more reactive around base 5666. This comparison suggests that the TR signal RNA structure can form at both 28°C and 37°C, whether this feature could promote TR in both mammalian and insect cells is currently unknown. The identified structures in this study agree with the previously determined structures [13].

**Figure 4.**
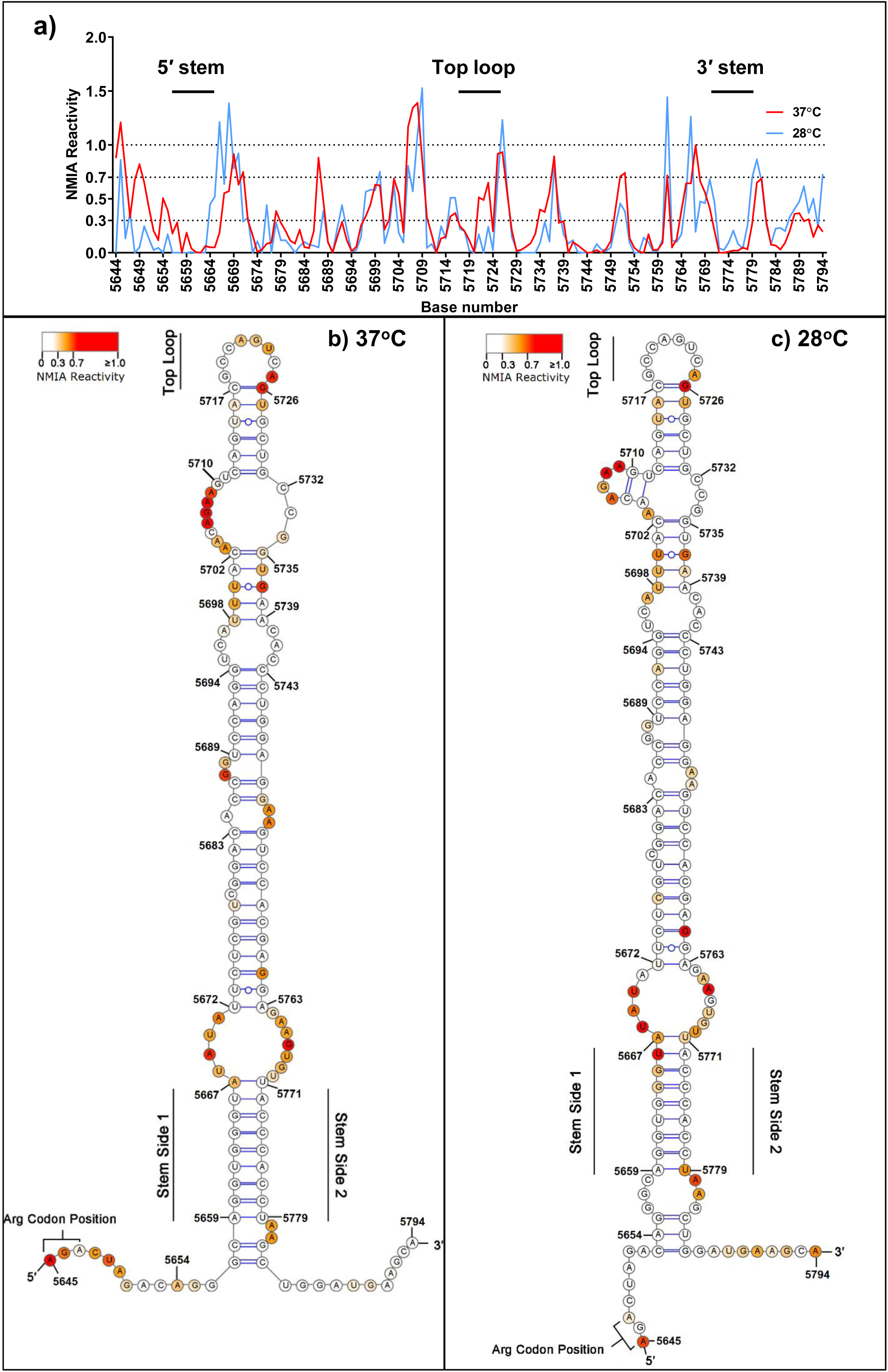
SHAPE analysis. (a) Graph of NMIA reactivity from nucleotide 5571 to 5721 (CHIKV nucleotide numbering from Genbank EU224268 (pCHIK-LR ic)) from RNA folded at either 37°C or 28°C. Numerical values presented in Supplementary Table S1. (b) Predicted structure between nucleotides 5572-5721 for RNA folded at 37°C, determined using RNAstructure software [20] constrained by the SHAPE data presented in (a). (c) As (b) but for RNA folded at 28°C.

**Figure 5.**
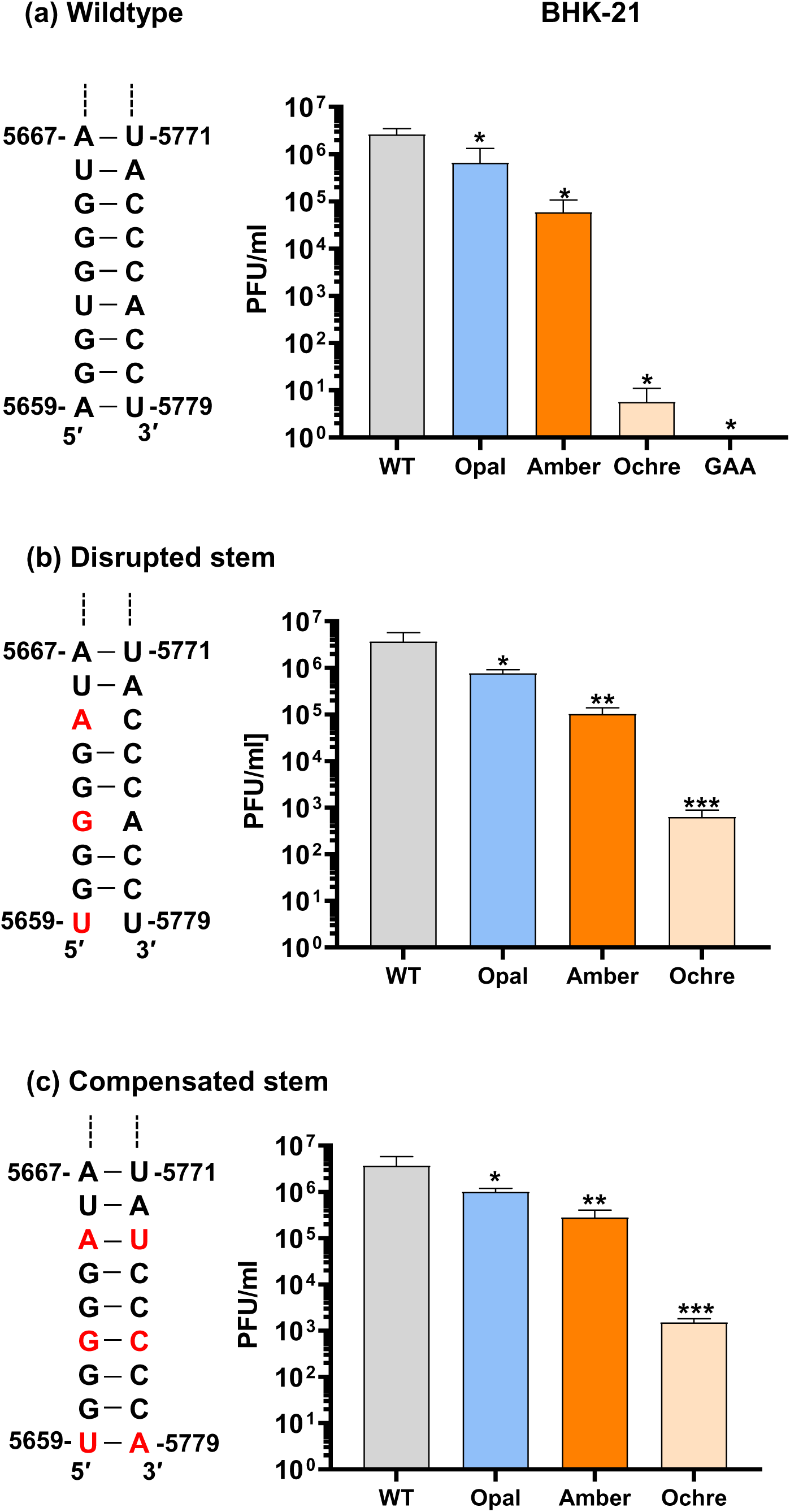
Infectivity analysis for stem mutants. (a) Sequence and structure of the base-paired region at the base of the stem from Fig. 4. Infectivity of wildtype and the 3 stop codon mutants following electroporation of *in vitro* transcribed RNA into BHK-21 cells. Note this is data from Fig. 3 (c), reproduced here for direct comparison. (b) Sequence and structure of the disrupted stem mutant (A5586U, U5589G, G5592A). Infectivity of the 3 stop codon mutants with this disrupted stem in BHK-21 cells. (c) Sequence and structure of the compensated stem mutant (mutations in (b) compensated by U5706A, A5703C, C5700U). Infectivity of the 3 stop codon mutants with this compensated stem in BHK-21 cells.

The short stem at the base of the RNA structure (residues 5659-5667 and 5771-5779) is highly conserved and stable at both 37°C and 28°C (Fig. 4). We therefore investigated the potential role of this stem in stop codon TR by mutagenesis. Three residues in the 5′ side of the stem were mutated to disrupt the structure, whilst maintaining the coding capacity. In a second construct, compensatory mutations were generated in the 3′ side to restore the stability of the stem. These mutants were generated in the context of the wildtype (arginine coding), opal, amber and ochre stop codon derivatives of the ICRES infectious clone (Fig. 3) and propagation of these mutants was evaluated following transfection into BHK-21 cells (Fig. 5). As shown in Fig. 5b, disruption of the stem had very little effect on the production of infectious CHIKV in BHK-21 cells, the pattern looking similar to the wildtype stem (Fig. 5a). Consequently when the stem structure was restored by the compensatory mutations (Fig. 5c) again the ability to produce infectious virus was similar to those seen in the context of the wildtype stem. We conclude that the stem structure is not absolutely required for stop codon TR in mammalian cells, and instead some other mechanism must be at play.

## Discussion

Misreading of a stop codon (TR) is estimated to occur spontaneously approximately 10^-5^ times per codon [21]. It seems unlikely that such a low frequency of TR would be compatible with the rapid and efficient characteristics of alphavirus replication as this would result in only one polymerase molecule for every 100,000 translation events. Having said that the fact that alphaviruses contain opal stop codons in preference to the other two stop codons, and the predominance of a C directly 3′ to the stop codon, correlates with the fact that an opal codon followed by a C is the most efficient context for TR [21]. Our data show that the presence of an opal stop codon has no detrimental effect on genome replication in any cells tested – human, murine or mosquito. We assume therefore that the frequency of TR at the opal stop codon must be consistent with the production of sufficient nsP4 RNA polymerase to allow genome replication to occur with the same efficiency as when an arginine is present. In other words the presence of the stop codon does not impair nsP4 translation and implies that alphaviruses possess a mechanism to potentiate TR at the opal stop codon.

It is intriguing that in the context of the SGR the amber stop codon is less well tolerated in mammalian cells but appears to be indistinguishable from an arginine codon in mosquito cells (Fig. 2). In addition replication of the ochre stop codon containing SGR was indistinguishable from the negative control GAA polymerase mutant in mammalian cells but was able to replicate almost as well as wildtype in mosquito cells. This suggests either that the mechanism and/or regulation of TR is distinct in mammalian and mosquito cells, or the low level of TR is sufficient to fully enable genome replication in mosquito cells. The latter seems unlikely as replication in C6/36 cells is as efficient as in mammalian cells.

We also note that in the context of infectious CHIKV the ochre stop codon resulted in a 1000-fold reduction in production of virus in C6/36 cells. Therefore, it may be that the reduced level of nsP4 translated following ochre stop codon TR impacts the production of the sgRNA encoding the structural proteins. In this context, our data agree with an early study which replaced the opal stop codon of the SINV with the other two stop codons; the production of the nsP34 polyprotein and the viral growth rate for the both the amber and ochre two stop codons were reduced with the ochre mutation exhibiting the largest decrease in viral growth rate of the two [14].

Unlike the results shown here, other studies have shown that the presence of the opal stop codon in alphaviruses enhances infection in the mosquito vector but not than the mammalian host. For example, in the case of O’Nyong Nyong virus (ONNV), increased infection of *Anopheles gambiae* mosquitoes was attributed to the effect of the opal stop codon on viral replication [22]. In another study passage of an ONNV strain containing the stop codon in mammalian cells (Vero) resulted in a switch of the stop codon to arginine [23], suggesting that the opal stop codon is selectively maintained in the mosquito vector but not the mammalian host. The inverse mutation of a La Réunion CHIKV strain possessing an opal stop codon into an arginine codon was shown to decrease viral titres produced in C6/36 cells (10^6^ vs 10^4^) [24]. The presence of an opal stop codon near the end of nsP3 must confer some advantage to the virus that is not apparent *in vitro*, either in mammalian or mosquito cells. It is possible that the opal stop codon plays a role in the complex natural life-cycle of the virus that involves replication and virus production in two unrelated species. However, the reason(s) why the presence of either an arginine or opal stop codon confers host specific advantages is currently unknown and will require further research to characterise.

SHAPE data confirmed the existence of a stable stem-loop structure in the CHIKV RNA from positions 5657-5783 (10 nucleotides 3′ of the stop codon) (Fig. 4). This structure was previously shown to enhance stop codon TR [12, 13]. Importantly, our data demonstrate that this structure is formed in the context of the full-length CHIKV genome and was stable at both 37°C and 28°C. The structure was largely similar at both temperatures although there were some differences at the base of the stem which was elongated at 28°C, extending from nucleotides 5653-5785. At the base of the stem is a conserved base-paired region that was observed at both temperatures. We hypothesised that the proximity of this region to the stop codon was consistent with a role in either promoting or enhancing TR so we tested this by disrupting the structure by mutation. However, surprisingly neither destabilizing mutations on one side of the stem, or additional compensatory mutations on the other side, had any effect on the production of infectious virus in BHK-21 cells. This was the case for both the arginine codon virus and the three stop codon versions. Therefore, we conclude that the conserved base-paired region at the base of the stem does not play a role in stop codon TR. It is possible that other structures within the stem-loop are more important for efficient TR, to evaluate this would require an extensive mutagenic study.

In conclusion our data demonstrate that the CHIKV lifecycle in either mammalian or mosquito cells is not dependent on the presence or absence of an opal stop codon near the end of nsP3. The other two stop codons are deleterious, particularly in mammalian cells. The biochemical mechanisms underpinning opal stop codon TR remain elusive and require further investigation. In this regard it will be of interest to determine if new pharmacological interventions that target TR [25] might be of efficacy as novel antiviral agents.

## Acknowledgements

RL was funded by a University of Leeds Research Scholarship. This work was supported by a Wellcome Investigator award (to MH: grant number 096670) and an MRC project grant (to AT: MR/N01054X/1). The funders had no role in study design, data collection and analysis, decision to publish, or preparation of the manuscript. We are grateful to Andres Merits (University of Tartu, Estonia) for the kind gift of CHIKV SGR and infectious clones.

## Materials and methods

### Tissue culture

Mammalian cells used in this study were murine C2C12 cells (muscle, myoblast), hamster BHK-21 cells (kidney, fibroblast), human Huh7 cells (liver, hepatocellular carcinoma); human SVG-A cells (brain, astroglia) and human RD cells (muscle, rhabdomyosarcoma), chosen as we had previously shown that these cells efficently supported CHIKV replication [16]. Mammalian cells were maintained in humidified incubator at 37°C with 5% CO_2_ in complete DMEM (Sigma) supplemented with 10% foetal calf serum (FBS) (Gibco) (or 20% FBS for C2C12 cells), 100 IU penicillin/ml and 100 µg/ml streptomycin (Gibco), 10% Non-essential amino acids (NEAA) (Thermo Fisher Scientific) and 10 mM HEPES (Thermo Fisher Scientific). The mosquito cell line C6/36 (*Ae. albopictus*, embryonic) used in this study has a mutant Dcr2 gene which results in a defective RNAi response [26]. These cells were maintained at 28°C.in Leibovitz’s L-15 Medium (Thermo Fisher Scientific) supplemented with 10% FBS, 100 IU penicillin/ml and 100 µg/ml streptomycin (Gibco), and 10% Tryptose phosphate broth (Thermo Fisher Scientific).

### Plasmid construction

The opal, amber and ochre stop codon were substituted into the CHIKV-Dluc-SGR [16, 27] using the Q5^®^ Site-Directed Mutagenesis Kit (NEB) as per manufacturers protocol. A two-step cloning strategy was used to incorporate the stop codons into the ICRES CHIKV infectious clone [15]. First, a 1017 bp GeneART (Thermo Scientific) synthesised DNA fragment matching the ICRES CHIKV sequence between the KflI (4683) and AgeI (5684) sites and containing two unique restriction sites (SpeI (5222) and SacI (5559)), was subcloned into the ICRES plasmid to generate an intermediate cloning plasmid. Fragments of nsP3 containing the stop codons from the CHIKV-Dluc-SGR constructs were then subcloned into SpeI-SacI digested ICRES plasmid. The stop codon ICRES CHIKV constructs with disrupted and compensated downstream RNA stem were generated using the Q5^®^ Site-Directed Mutagenesis Kit as described previously. The nsP4 GAA mutant CHIKV-Dluc-SGR and ICRES CHIKV infectious clone was generated previously in our laboratory [28].

### Transfection and Dual-luciferase assay

Purified linearised DNA was used as a template for capped *in vitro* RNA transcription using the mMESSAGE mMACHINE^®^ SP6 kit (Thermo Fisher Scientific), RNA was purified using lithium chloride precipitation and resuspended in RNAse-free H_2_O. Cells were seeded at 1 x 10^5^ cells per well in a 24 well plate for transfection using lipofectamine 2000 (Thermo Fisher Scientific) as per manufacturers protocol. Each well was transfected with 250 ng of RNA, using 1 µl of lipofectamine and 100 µl of opti-MEM™. Cells were lysed at the indicated timepoints using passive lysis buffer (Promega) and analysed using the Dual-luciferase^®^ Reporter Assay System (Promega), a FLUOstar Omega Microplate reader and Optima software (BMG LABTECH).

### Genome alignment

Genome sequences of CHIKV and related alphaviruses were aligned using the MAFFT Multiple sequence alignment software [29] and the resultant alignment was viewed using the Jalview software [30].

### Generation of infectious ICRES CHIKV

Capped ICRES RNA was generated by *in vitro* transcription using the mMESSAGE mMACHINE^®^ SP6 kit. For virus propagation in BHK-21 cells, 10^6^ cells in 400 µl medium were electroporated with 1 µg of CHIKV RNA using a square wave electroporation protocol (260 V, 25 ms, 1 pulse in a 4 mm cuvette). The electroporated cells were resuspended in 10 ml of complete media and incubated at 37°C. For propagation in C6/36 cells, 4 x 10^5^ cells were seeded in each well of a 6 well plate and transfected with 1 µg of ICRES CHIKV RNA, using 5 µl of lipofectamine 2000 and 500 µl of Opti-MEM. The transfection mixture was removed after 4 h, cells were washed, and incubated in complete Leibovitz’s L-15 Medium at 28°C for 48 h. Supernatants were collected at 48 h post transfection, the virus stocks were clarified (1000 x g, 5 min, room temperature) before aliquoting and storage at – 80°C.

### Titration of CHIKV by plaque assay

For plaque assay BHK-21 cells were seeded in either 6 or 12 well plates at 4 x 10^5^ or 2 x 10^5^ cells per well respectively. Virus dilutions were added and incubated at 37°C for 1 h. The inoculum was removed, cells washed with 1 ml of PBS and overlaid with 1.6% methyl cellulose (MC) in complete media. After 48 h incubation at 37°C the overlay was removed, and cells were fixed with 4% paraformaldehyde (PFA) for 30 min, stained with 0.5% (w/v) crystal violet solution and washed with water until plaques were visible.

### 2**^′^**hydroxyl acylation analysed by primer extension (SHAPE)

SHAPE analysis was performed as previously described [31]. Briefly, 10 pmol of CHIKV genomic RNA was denatured at 95°C for 3 min, incubated for 3 min on ice before addition of 6 µl of folding buffer (330 mM HEPES (pH 8.0), 20 mM MgCl_2_, 330 mM NaCl) and allowed to refold at either 37°C or 28°C for 20 min. Samples were then divided into positive and negative reactions and incubated with either 1 µl of 100 mM *N*-methylisatoic anhydride (NMIA) (positive) or 1 µl of DMSO (negative) for 45 min at 37°C or 28°C. Each reaction was terminated by ethanol and re-suspended in 10 µl 0.5 X TE containing RNAsecure (Thermo Fisher Scientific). For primer extension of the positive and negative reactions, 5 µl of resuspended RNA was incubated with 1 µl of 10 mM 5′ FAM labelled fluorescent oligonucleotide primer (ICRES nt positions 318–337) (Sigma-Aldrich) and 6 µl ddH2O at 85°C for 1 min, 60°C for 10 min and 30°C for 10 min. A master mix of 4 µl Superscript III reverse transcriptase buffer, 1 µl 100 mM DTT, 0.5 µl 100 mM dNTPs, 0.5 µl RNAseOUT, 1 µl ddH2O and 1 µl Superscript III reverse transcriptase (Thermo Fisher Scientific) was added to each reaction, which were incubated for 30 min at 55°C.

For generating SHAPE sequencing ladders, 6 pmol of *in vitro* transcribed RNA in 7.5 µl 0.5×TE buffer, 1 µl of 10 mM 5′ HEX labelled oligonucleotide primer (Sigma-Aldrich) and 2 µl ddH2O was incubated at 85°C for 1 min, 60°C for 10 min and 30°C for 10 min. A master mix of 4 µl Superscript III reverse transcriptase buffer, 1 µl 100 mM DTT, 0.5 µl 100 mM dNTPs, 0.5 µl RNAseOUT, 2 µl ddGTP and 1 µl Superscript III reverse transcriptase was added before incubation for 30 min at 55°C. Following incubation, all extensions were heated at 95°C for 3 min with 1 µl 4 M NaOH before cooling on ice with 2 µl 2M HCl for 2 min. Following ethanol precipitation cDNA was resuspended in 40 µl deionized formamide and pooled with 20 µl of SHAPE sequencing ladder.

### SHAPE data analysis and RNA structure prediction

Fragment size analysis of SHAPE extension products was conducted by capillary electrophoresis (DNA Sequencing and Services; part of the MRC-PPU Reagents and Services Facility, College of Life Sciences, University of Dundee, Scotland). SHAPE data was processed and normalized using the QuSHAPE software with default settings [19]. As previously published, based on an average of at least three independent biological repeats, nucleotides with normalized SHAPE reactivities 0– 0.3, 0.3–0.7 and >0.7 were taken to be unreactive, moderately reactive, and highly reactive respectively. *In silico* thermodynamic RNA structure and free energy predictions were carried out using UNAFOLD version 2.3 at 28°C and 37°C [32]. Normalized SHAPE reactivities were used as constraints to generate a thermodynamic RNA structure model using the RNA structure software [20, 33]. RNA structures were visualized and overlaid with normalized SHAPE reactivities using the VARNA software [34].

