## Supplementary Table 1 for "A structural and functional analysis of opal stop codon translational readthrough during Chikungunya Virus replication"

| Base position | 37°C | 28°C |
|---|---|---|
| **5644** | 0.8775 | 0 |
| 5645 | 1.2075 | 0.86 |
| **5646** | 0.7675 | 0.135 |
| 5647 | 0.33 | 0.37 |
| **5648** | 0.675 | 0 |
| 5649 | 0.82 | 0.05 |
| **5650** | 0.6525 | 0.245 |
| 5651 | 0.4175 | 0.13 |
| **5652** | 0.2025 | 0.025 |
| 5653 | 0.1175 | 0.045 |
| **5654** | 0.505 | 0 |
| 5655 | 0.3925 | 0.17 |
| **5656** | 0.18 | 0 |
| 5657 | 0.28 | 0 |
| **5658** | 0 | 0 |
| 5659 | 0.185 | 0 |
| **5660** | 0.04 | 0.005 |
| 5661 | 0.005 | 0 |
| **5662** | 0 | 0 |
| 5663 | 0.0625 | 0 |
| **5664** | 0.05 | 0.45 |
| 5665 | 0.05 | 0.52 |
| **5666** | 0.185 | 1.21 |
| 5667 | 0.5525 | 0.64 |
| **5668** | 0.57 | 1.385 |
| 5669 | 0.9125 | 0.785 |
| **5670** | 0.6325 | 0.92 |
| 5671 | 0.7475 | 0.29 |
| **5672** | 0.3 | 0.325 |
| 5673 | 0.1625 | 0 |
| **5674** | 0.0075 | 0.105 |
| 5675 | 0.03 | 0 |
| **5676** | 0.0825 | 0.44 |
| 5677 | 0.095 | 0 |
| **5678** | 0.385 | 0.28 |
| 5679 | 0.2775 | 0.115 |
| **5680** | 0.2025 | 0.115 |
| 5681 | 0.1125 | 0.065 |
| **5682** | 0.085 | 0 |
| 5683 | 0.2125 | 0.06 |
| **5684** | 0.0525 | 0.1 |
| 5685 | 0.0425 | 0.075 |
| **5686** | 0.2 | 0.065 |
| 5687 | 0.88 | 0.05 |
| **5688** | 0.49 | 0.385 |
| 5689 | 0.1 | 0.095 |
| **5690** | 0.0025 | 0 |
| 5691 | 0.15 | 0.23 |

| | | |
|---|---|---|
| **5692** | 0.285 | 0.44 |
| 5693 | 0.1175 | 0.215 |
| **5694** | 0.015 | 0.01 |
| 5695 | 0.055 | 0.025 |
| **5696** | 0.2525 | 0.145 |
| 5697 | 0.3375 | 0.565 |
| **5698** | 0.4475 | 0.585 |
| 5699 | 0.63 | 0.58 |
| **5700** | 0.625 | 0.75 |
| 5701 | 0.2175 | 0.09 |
| **5702** | 0.31 | 0.24 |
| 5703 | 0.685 | 0.625 |
| **5704** | 0.5475 | 0.185 |
| 5705 | 0.1875 | 0.095 |
| **5706** | 1.1675 | 0.805 |
| 5707 | 1.345 | 0.57 |
| **5708** | 1.39 | 1.095 |
| 5709 | 0.8625 | 1.525 |
| **5710** | 0.3275 | 0.045 |
| 5711 | 0.17 | 0.08 |
| **5712** | 0 | 0 |
| 5713 | 0.1325 | 0.245 |
| **5714** | 0.1475 | 0.135 |
| 5715 | 0.3375 | 0.51 |
| **5716** | 0.3675 | 0.51 |
| 5717 | 0.2475 | 0.22 |
| **5718** | 0.2 | 0.2 |
| 5719 | 0.115 | 0.02 |
| **5720** | 0.0075 | 0 |
| 5721 | 0.5125 | 0.17 |
| **5722** | 0.49 | 0 |
| 5723 | 0.6475 | 0.24 |
| **5724** | 0.2 | 0.025 |
| 5725 | 0.9175 | 0.66 |
| **5726** | 0.9325 | 1.23 |
| 5727 | 0.5625 | 0.6 |
| **5728** | 0.21 | 0.15 |
| 5729 | 0.0175 | 0.025 |
| **5730** | 0.0275 | 0 |
| 5731 | 0.055 | 0 |
| **5732** | 0.0875 | 0 |
| 5733 | 0.175 | 0 |
| **5734** | 0.4 | 0.09 |
| 5735 | 0.3775 | 0.285 |
| **5736** | 0.56 | 0.105 |
| 5737 | 0.8925 | 0.79 |
| **5738** | 0.2775 | 0.42 |
| 5739 | 0.2925 | 0.17 |
| **5740** | 0 | 0.085 |
| 5741 | 0.0975 | 0.115 |

| | | |
|---|---|---|
| **5742** | 0.07 | 0.005 |
| 5743 | 0.035 | 0 |
| **5744** | 0 | 0 |
| 5745 | 0.0025 | 0 |
| **5746** | 0.025 | 0.08 |
| 5747 | 0 | 0.03 |
| **5748** | 0.035 | 0.015 |
| 5749 | 0.11 | 0.05 |
| **5750** | 0.4075 | 0.255 |
| 5751 | 0.7075 | 0.455 |
| **5752** | 0.74 | 0.385 |
| 5753 | 0.1325 | 0.09 |
| **5754** | 0.045 | 0 |
| 5755 | 0.01 | 0 |
| **5756** | 0 | 0.14 |
| 5757 | 0.17 | 0.1 |
| **5758** | 0.025 | 0.01 |
| 5759 | 0.03 | 0.02 |
| **5760** | 0.0925 | 0.125 |
| 5761 | 0.715 | 1.44 |
| **5762** | 0.05 | 0.115 |
| 5763 | 0.125 | 0.1 |
| **5764** | 0.3925 | 0.17 |
| 5765 | 0.6425 | 0.525 |
| **5766** | 0.6425 | 1.26 |
| 5767 | 0.995 | 0.195 |
| **5768** | 0.66 | 0.475 |
| 5769 | 0.56 | 0.46 |
| **5770** | 0.4125 | 0.68 |
| 5771 | 0.2025 | 0.465 |
| **5772** | 0 | 0.04 |
| 5773 | 0 | 0.025 |
| **5774** | 0.01 | 0.095 |
| 5775 | 0.02 | 0.205 |
| **5776** | 0.02 | 0.08 |
| 5777 | 0.0475 | 0.04 |
| **5778** | 0.0325 | 0.25 |
| 5779 | 0.2675 | 0.7 |
| **5780** | 0.65 | 0.865 |
| 5781 | 0.6875 | 0.66 |
| **5782** | 0.29 | 0.265 |
| 5783 | 0.075 | 0.08 |
| **5784** | 0.0675 | 0.225 |
| 5785 | 0.03 | 0.23 |
| **5786** | 0.0925 | 0.2 |
| 5787 | 0.205 | 0.335 |
| **5788** | 0.36 | 0.405 |
| 5789 | 0.3675 | 0.475 |
| **5790** | 0.2925 | 0.615 |
| 5791 | 0.31 | 0.37 |

| | | |
|---|---|---|
| **5792** | 0.146667 | 0.5 |
| 5793 | 0.255 | 0.24 |
| **5794** | 0.195 | 0.73 |

**Supplementary Table S1.** Normalized SHAPE reactivities for CHIKV nucleotides 5644 to 5799 from full-length genomic RNA folded at either 37oC or 28oC, n=3.
